# Donor-Targeted Anti-HLA-A2 Antibody Shows Nanogram-Level Efficacy in a Novel GVHD Mouse Model

**DOI:** 10.1101/2025.07.16.665074

**Authors:** Kouta Niizuma, Alyssa H. Chang, Jeewon S. Kim, Stephan Busque, Ravindra Majeti, Hiromitsu Nakauchi, Yusuke Nakauchi

## Abstract

Acute graft-versus-host disease (GVHD) remains a life-threatening complication of allogeneic hematopoietic stem cell transplantation (allo-HSCT), with limited treatment options for steroid-refractory cases. Current therapies broadly suppress immune responses, increasing infection risk and impairing graft function. We hypothesized that selective depletion of donor immune cells through HLA allele-specific targeting could offer a safer and more precise alternative. To test this, we developed a monoclonal antibody (AN7) against HLA-A2 by immunizing HLA-A24 transgenic mice with recombinant HLA-A2 tetramers, followed by hybridoma screening. AN7 specifically recognized HLA-A2 and HLA-A68, and was reformatted into murine IgG2a and human chimeric IgG1 recombinant antibodies. These retained binding specificity and mediated robust complement-dependent cytotoxicity (CDC), antibody-dependent cellular cytotoxicity (ADCC), and antibody-dependent cellular phagocytosis (ADCP) against HLA-A2+ target cells. To evaluate therapeutic potential, we established a fully MHC-mismatched allo-HSCT model using HLA-A2 transgenic donor bone marrow in lethally irradiated BALB/c recipients. A single ultra-low dose of AN7 (1-100 ng) administered post-transplant significantly improved survival and reduced GVHD while preserving donor hematopoiesis. These findings highlight the potential of allele-specific antibody-mediated clearance as a mechanistically distinct approach to donor immune modulation in GVHD.

## TO THE EDITOR

Acute graft-versus-host disease (GVHD) remains a major barrier to successful allogeneic hematopoietic stem cell transplantation (allo-HSCT), with severe cases often being fatal despite aggressive immunosuppression. While novel agents continue to emerge, severe or steroid-refractory GVHD remains associated with poor outcomes and limited treatment efficacy^1-4^. GVHD following solid organ transplantation (SOT), though rarer, is equally challenging and lacks effective targeted therapies^5,6^.

Although current acute GVHD treatment relies on a stepwise intensification of immunosuppression, starting with immunosuppressants like steroids, followed by JAK inhibitors and antibody therapies, we hypothesized that selective depletion of donor alloreactive immune cells via HLA allele-specific targeting could offer a precise and clinically actionable therapeutic approach.

In our prior study, we developed a murine monoclonal antibody against HLA-A2 and demonstrated its capacity to attenuate GVHD in a xenogeneic peripheral blood mononuclear cells (PBMCs) transfer model (Human HLA-A2^+^ PBMCs were transferred to immunodeficient mice without any conditioning)^7^. However, that model lacked HSCT and did not allow assessment of dose sensitivity or therapeutic selectivity—critical factors for translational application.

The clinical potential of donor-directed serotherapy was recently underscored by a compelling case in which transfusion of plasma containing high titers of donor-allele-specific (anti-HLA-A2) antibodies led to complete resolution of steroid-refractory GVHD following kidney transplantation^8^. Although polyclonal in nature and not dose-optimized, this case illustrates that targeted depletion of mismatched donor immune components can be therapeutically effective, supporting the rationale for developing allele-specific monoclonal antibodies with defined pharmacologic properties.

Motivated by these insights, we developed a new monoclonal antibody with allele-specific binding to test whether donor-targeted depletion could provide a safe, selective, and clinically scalable strategy for GVHD treatment.

Here, we report the development of AN7, a new anti-HLA-A2 monoclonal antibody with improved translational potential. AN7 was generated by immunizing HLA-A24 transgenic mice with recombinant HLA-A2 tetramer: a strategy designed to preferentially elicit allele-specific immune responses while minimizing cross-reactivity to conserved HLA epitopes in immunized animals^9^. Hybridoma clones were screened using HLA-coated beads and HLA-A2□PBMCs, and a monoclonal antibody (clone name; AN7) was selected based on its potent and selective binding to HLA-A2 and the closely related allele HLA-A68^10^ (Figure 1A, B). Isotyping via a strip assay revealed that AN7 is a mouse IgG2aκ antibody (Figure 1C).

**Figure 1.**
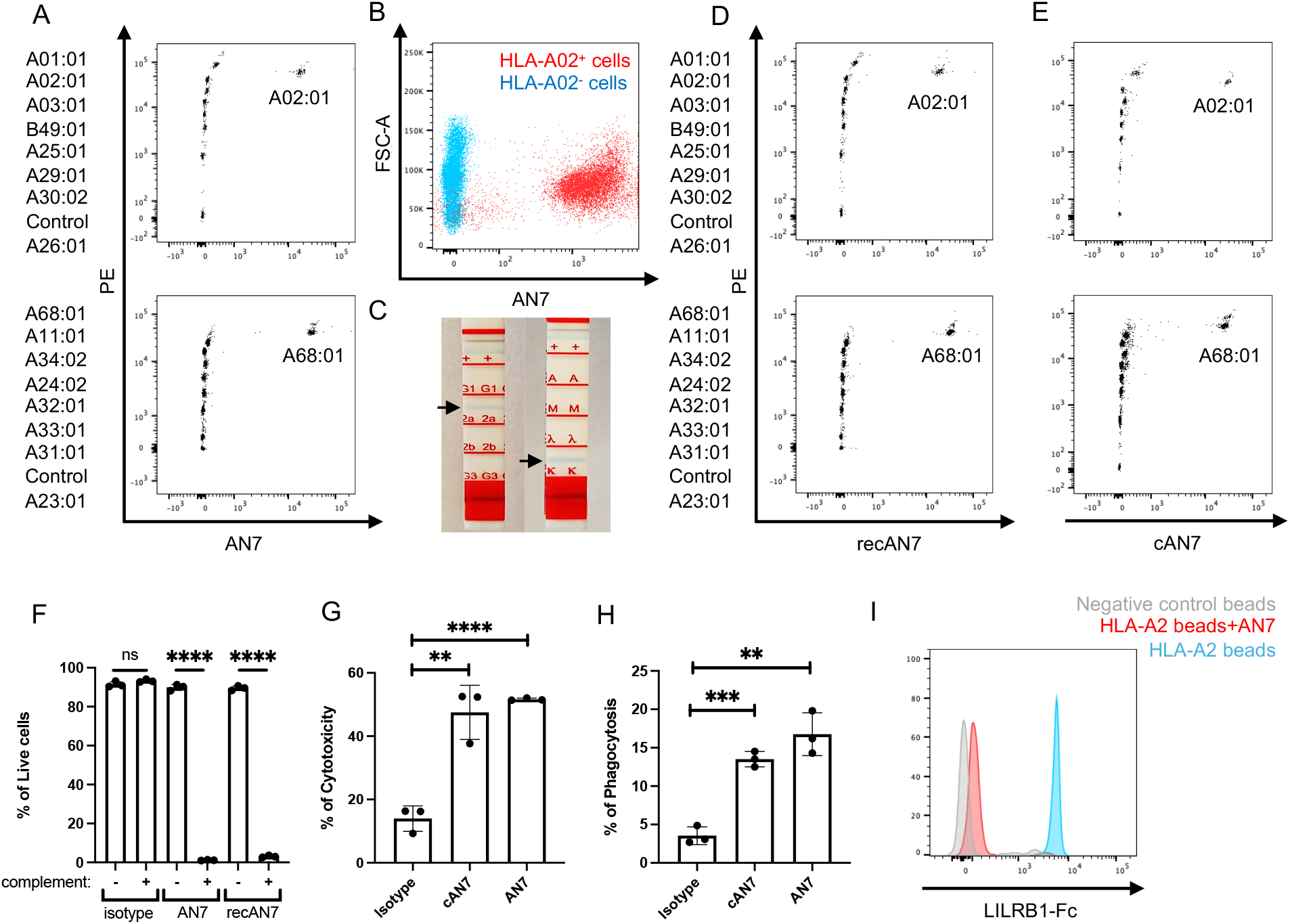
Generation and characterization of the anti-HLA-A2 monoclonal antibody AN7. A. Flow cytometric analysis of HLA allele reactivity using LABScreen Single Antigen Class I beads incubated with hybridoma-derived AN7. B. Flow cytometric analysis of AN7 binding to HLA-A2□ and HLA-A2□ PBMCs from healthy donors. C. Isotyping of AN7 using a strip assay. D. Flow cytometric analysis of HLA allele reactivity using LABScreen Single Antigen Class I beads incubated with recombinant AN7 (mouse IgG2aκ). E. Flow cytometric analysis of HLA allele reactivity using LABScreen Single Antigen Class I beads incubated with chimeric AN7 (human IgG1κ). F. Complement-dependent cytotoxicity (CDC) assay using human PBMC-derived HLA-A2□ cells. Cells were incubated with monoclonal antibodies (AN7, recAN7, or isotype control) in the presence or absence of baby rabbit complement. Cell viability was assessed by flow cytometry. G. Antibody-dependent cellular cytotoxicity (ADCC) assay using Calcein-AM-labeled HLA-A2□ NALM6 target cells and HLA-A2□ NK cells at a 5:1 E:T ratio. Cells were treated with AN7, cAN7, or isotype control (0.5 µg/well). Specific lysis was quantified by fluorescence measurement of released Calcein. H. Antibody-Dependent Cellular Phagocytosis (ADCP) Assay: HLA-A2^-^ monocyte-derived macrophages were co-cultured with GFP-expressing HLA-A2^+^ NALM6 target cells at a 1:2 effector-to-target (E:T) ratio in the presence of isotype control antibody, cAN7, or AN7. Phagocytic activity was quantified by flow cytometry as the percentage of macrophages containing GFP^+^ target cells. Data represent mean ± SEM from three independent experiments (*p < 0.05, **p < 0.01, ***p < 0.001). I. LILRB1 binding assay. HLA-A2-coated or negative control beads were incubated with or without AN7 prior to the addition of LILRB1-Fc. Binding was analyzed by flow cytometry using a FITC-conjugated anti-human IgG Fc antibody.

The variable regions of AN7 were sequenced to generate recombinant formats: a murine IgG2aκ (recombinant AN7; recAN7) and a human chimeric IgG1κ version (cAN7). Flow cytometry–based binding assays confirmed that recAN7 and cAN7 antibodies preserved the HLA-A2/A68 specificity (Figure 1D, E).

Functionally, AN7 and recAN7 exhibited strong complement-dependent cytotoxicity (CDC) against HLA-A2□ PBMCs (Figure 1F). Both AN7 and cAN7 also mediated significant antibody-dependent cellular cytotoxicity (ADCC) against HLA-A2□ PBMCs by HLA-A2^-^ NK cells and robust antibody-dependent cellular phagocytosis (ADCP) against HLA-A2□ target cell lines by HLA-A2^-^ macrophages (Figure 1G, H).

Since HLA class I molecules are known ligands for LILRB1, an inhibitory receptor expressed on myeloid and NK cells, we hypothesized that AN7 might also function as a checkpoint inhibitor^11^. In a competitive binding assay using recombinant LILRB1-Fc, pre-incubation of HLA-A2 beads with AN7 markedly reduced subsequent LILRB1-Fc binding (Figure 1I). These results suggest that AN7 not only triggers direct effector responses via CDC, ADCC, and ADCP but may also promote immune activation by disrupting LILRB1-mediated inhibitory signaling.

Next, we evaluated the therapeutic efficacy of AN7 in two preclinical models of GVHD. We first tested AN7 in a widely used xenogeneic PBMC transfer model, in which 10 million HLA-A2□PBMCs are transferred into immunodeficient mice (NSG) on day 0. In this setting, human T cells rapidly expand and attack murine tissues, resulting in lethal GVHD^12^. A single dose of 100 μg AN7 or recAN7 on day 2 significantly improved survival compared to isotype-treated controls (Figure 2A).

**Figure 2.**
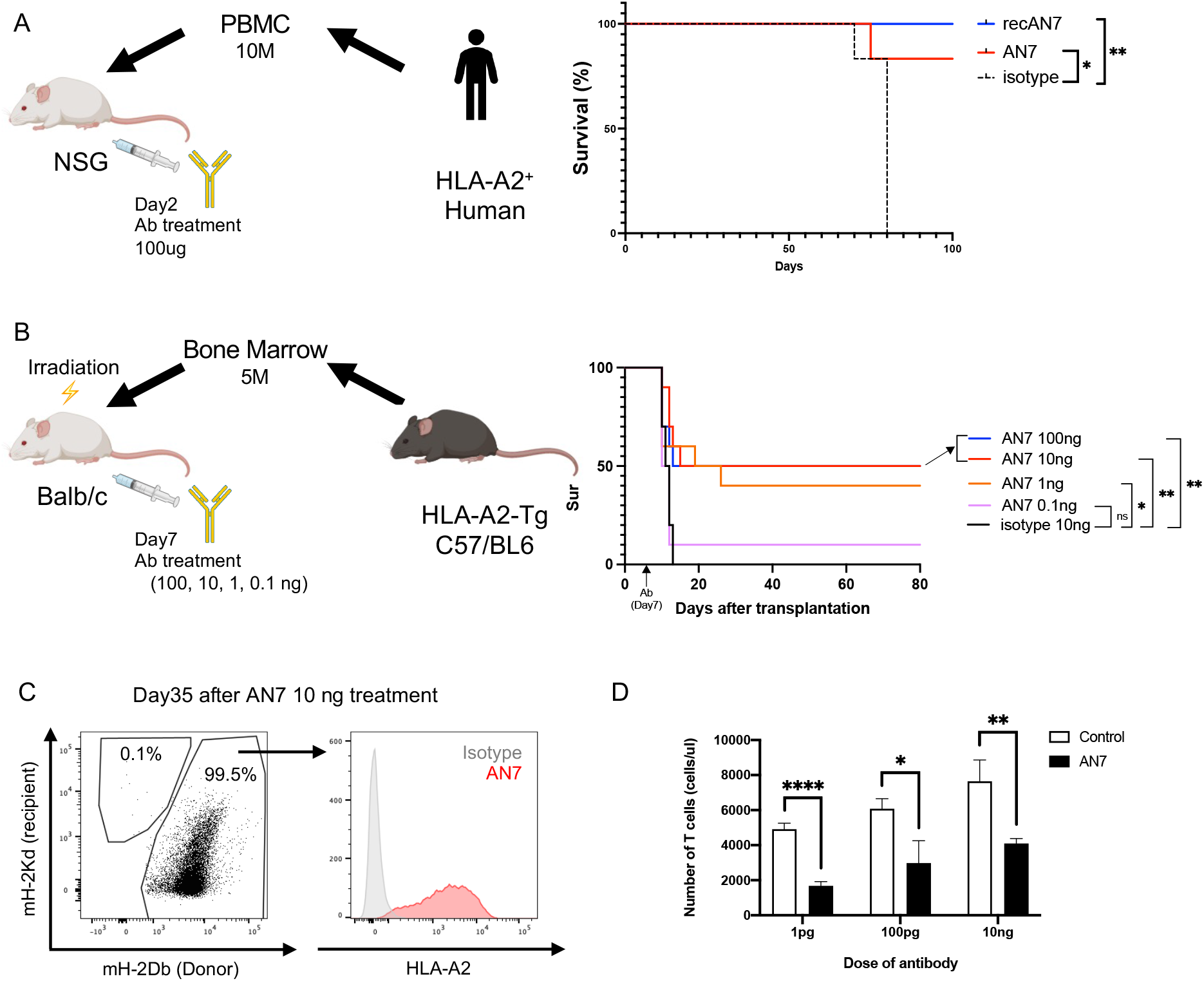
AN7 Ameliorates GVHD in two Preclinical Transplant Models. A. Schematic and survival analysis of xenogeneic GVHD model in NSG mice. HLA-A2□ human PBMCs (1□×□10□ cells) were injected intravenously into non-irradiated NSG mice. On day 2 post-transplantation, mice were treated intraperitoneally with 100□µg of AN7, recombinant AN7, or isotype control. Survival was monitored and plotted as Kaplan–Meier curves (n = 6 per group). B. Schematic and survival analysis of a fully allogeneic GVHD model. Lethally irradiated BALB/c mice received 5 × 10□ bone marrow cells from HLA-A2 transgenic C57BL/6 donors. On day 7 post-transplantation, mice were treated with a single dose of AN7 (0.1□ng, 1□ng, 10□ng, or 100□ng,) or mouse IgG isotype control (10 ng). Survival was monitored and plotted as Kaplan– Meier curves. Statistical comparisons were performed as indicated. ns = not significant; *P < 0.05; **P < 0.01. C. Representative flow cytometry data using peripheral blood cells from a BALB/c recipient that received a bone marrow transplant from HLA-A2 transgenic C57BL/6 donors, followed by treatment with 10□ng of AN7. Analysis was performed 35 days after antibody administration. Anti-mH-2Kd and anti-mH-2Dd antibodies were used to detect recipient and donor chimerism, respectively. D. Mixed lymphocyte reaction (MLR) assay. Splenocytes from HLA-A2 transgenic C57BL/6 mice (responders) and mitomycin C–treated BALB/c splenocytes (stimulators) were co-cultured at a 1:1 ratio in the presence of AN7 (1□pg, 100□pg, 10□ng) or isotype control antibody. After 72 hours, T cell counts were determined by flow cytometry using counting beads. Data represent mean ± SD from triplicate wells (n = 3). Statistical significance: *P < 0.05, **P < 0.01, ***P < 0.001, ****P < 0.0001 (paired t-test).

While this model demonstrates the capacity of AN7 to deplete donor immune cells and mitigate GVHD, it lacks HSCT and thus cannot address a critical translational question: whether AN7 can be dosed to eliminate pathogenic donor cells while sparing essential HSCs. In clinical settings, GVHD prevention alone is not sufficient; therapies must spare donor-derived hematopoietic stem cells or organ function to ensure durable engraftment and transplant success^13^. To address this limitation, we established a new immunocompetent allo-GVHD model using lethally irradiated BALB/c mice transplanted with 5 million bone marrow cells from HLA-A2 transgenic C57BL/6 donors^14^. This fully MHC-mismatched combination induces robust and rapidly fatal GVHD. In addition, because most HSCs in the host BM are of donor origin, we can assess the effect of AN7 on donor HSCs.

To determine the therapeutic window of AN7, we administered a single dose at varying concentrations (0.1, 1, 10, 100□ng) on day 7 post-transplant, alongside a 10□ng isotype control. The rationale for evaluating such ultra-low doses was based on clinical evidence suggesting that donor-specific antibodies can mediate therapeutic effects at concentrations estimated ≤6 μg/kg^8^. Strikingly, a single administration of AN7 at 1, 10, or 100□ng conferred long-term survival in approximately half of the mice, whereas all animals in the isotype control and 0.1□ng groups succumbed within two weeks (Figure 2B). These findings define an effective and well-tolerated dose range of 1–100□ng for AN7 in the new GVHD model.

Importantly, all survivors exhibited successful donor hematopoietic reconstitution, indicating that AN7 selectively eliminates alloreactive immune cells while sparing donor hematopoietic stem cells (HSCs) (Figure 2C). This selectivity is consistent with known HSC biology; HSCs residing in the BM niche are reported to exhibit resistance to apoptosis and immune-mediated attacks.^15,16^

To further validate the donor-specific immunosuppressive activity of AN7, we conducted an *in vitro* mixed lymphocyte reaction (MLR), using mitomycin C-treated BALB/c splenocytes as stimulators and splenocytes from HLA-A2 transgenic C57BL/6 mice as responders. Treatment with AN7 at picogram-range concentrations (1□pg to 10□ng) significantly reduced the expansion of T cells (Figure 2D), reinforcing the specificity and potency of AN7-mediated immunosuppression.

Our findings establish AN7 as a promising donor-targeted therapeutic with unprecedented efficacy at nanogram-level doses in the GVHD models. In contrast to conventional strategies that broadly suppress immune function, our strategy selectively targets a mismatched donor allele, enabling precise depletion of alloreactive cells while sparing stem cells. Remarkably, robust *in vivo* activity was achieved with as little as 1–100□ng per mouse (0.05-5□μg/kg), a 1,000- to 100,000-fold reduction compared to typical therapeutic antibody doses used in clinical settings. This ultra-low dose requirement implies not only a favorable safety margin but also offers clear advantages in terms of manufacturing efficiency, cost-effectiveness, and clinical scalability.

HLA-A2 is among the most frequently mismatched alleles in allogeneic transplantation, owing to its high prevalence across global populations^17,18^. AN7 exemplifies a novel class of allele-specific antibodies capable of selectively eliminating donor alloreactive immune cells. The use of a human chimeric antibody format enhances translational relevance and supports the clinical applicability of this approach. Supported by data from a clinically relevant GVHD model, our study lays the foundation for next-generation immunotherapies with improved specificity and reduced off-target toxicity. This approach can be extended to other common HLA alleles, enabling broadly applicable precision immunotherapy.

## Materials and Methods

### Cells

Healthy donor human peripheral blood cells (PBMC) were obtained from the Stanford Blood Center. Mononuclear cells from peripheral blood and cord blood were isolated using Ficoll-Paque (Fisher Scientific, # 45-001-750) separation. The NALM6 cell line was cultured in RPMI-1640 medium supplemented with 10% fetal bovine serum (FBS). HLA typing of all cell types was performed by staining with either our in-house anti-HLA-A2 antibody (AN7) or a commercially available anti-HLA-A2 antibody (clone BB7.2), followed by flow cytometric analysis to confirm HLA expression.

### Mice

NOD.Cg-Prkdc^scid Il2rg^tm1Wjl/SzJ (NSG), HLA-A2 transgenic (RRID:IMSR_JAX:003475), and BALB/c mice (RRID:IMSR_JAX:000651) were obtained from Jackson Laboratory. All mice were bred and housed in the animal research facility at Stanford University. The animals were maintained under specific pathogen-free conditions with ad libitum access to food and water, in accordance with Stanford University’s Administrative Panel on Laboratory Animal Care (APLAC) guidelines and the Guide for the Care and Use of Laboratory Animals. All procedures were approved by the Stanford University APLAC prior to initiation, and animal care was provided in compliance with institutional, state, and federal regulations.

### Isotyping assay

Isotype determination was performed using the Mouse Monoclonal Antibody Isotyping Kit (Bio-Rad, # MCA871K) following the manufacturer’s instructions. The assay employs a lateral flow strip format pre-coated with capture reagents specific for mouse immunoglobulin isotypes and light chains. Purified AN7 antibody was applied directly to the strip, and isotype was determined by visual inspection within 10 minutes.

### Antibody production and purification

Hybridoma cells were injected intraperitoneally into NSG mice. Ascites fluid was collected 10-14 days after the injection. Monoclonal antibodies were purified from the ascites using Protein A/G affinity chromatography according to standard protocols. Purified antibodies were quantified and stored at −80□until use.

### Flow cytometry using HLA-beads

The specificity of each anti-HLA antibody was determined using HLA-beads, each coated with individual HLA antigens (ThermoFisher, FlowPRA HLA Class I Screening Test). The binding of anti-HLA antibodies to specific HLA antigens was assessed by flow cytometry. All procedures were performed according to the manufacturer’s instructions provided with the product.

### Complement-Dependent Cytotoxicity (CDC) assay

Target cells were seeded at a density of 1 × 10^5^ cells per well in 200 μL of medium in 96-well flat-bottom plates. Antibodies were diluted to the indicated concentrations in medium and added to the appropriate wells (0.6 μg/well). The plate was incubated at room temperature for 5 minutes to allow antibody binding. Subsequently, baby rabbit complement (Cedarlane, #CL3441-S50-R) was added to each well at a final concentration of 10%. Plates were then incubated at 37°C for 2 hours to enable complement-mediated cytotoxicity. After incubation, cell viability was assessed by flow cytometry using propidium iodide (PI) staining to distinguish live and dead cells. Unless otherwise specified, all cell culture reagents consisted of RPMI 1640 supplemented with 10% fetal bovine serum (FBS) and penicillin/streptomycin.

### Antibody-Dependent Cellular Cytotoxicity (ADCC) assay

The ADCC assay was conducted using HLA-A2-negative natural killer (NK) cells isolated from human PBMCs with the EasySep CD56 Positive Selection Kit (Stemcell Technologies, #17855). The isolated NK cells were cultured overnight in RPMI 1640 medium supplemented with 10% FBS and 100 U/mL recombinant human IL-2 (PeproTech, #200-02), then resuspended in HBSS + 10% FBS. HLA-A2-positive target cells (NALM6) were labeled with 5 µM Calcein-AM (Thermo Fisher, #C3100MP) for 30 minutes at 37°C, washed, and then resuspended in HBSS + 10% FBS. Antibodies (mAN7, chimeric AN7 (cAN7), and isotype antibody as negative control) were evaluated at 0.5 µg/well alongside an isotype control. The labeled target cells (4,000 cells/well) and activated NK cells (20,000 cells/well) were co-cultured at a 1:5 target-to-effector (T:E) ratio in V-bottom plates for 2 hours at 37°C. Promega lysis buffer (Promega, #G182A) served as the maximal lysis control. Supernatant fluorescence (490/520 nm) was measured using a SpectraMax M3 plate reader. Percent specific lysis was calculated as: [(test RFU - mean background RFU)/(mean maximal RFU - mean background RFU)] * 100, where background is effector cells + target cells without antibody. Maximal lysis is defined as effector cells + target cells with lysis buffer (Promega, #G182A).

### Antibody-Dependent Cellular Phagocytosis (ADCP) assay

ADCP was evaluated using HLA-A2-negative monocyte-derived macrophages as effector cells. Human PBMCs were cultured in IMDM supplemented with 10% human AB serum, penicillin/streptomycin, and Glutamax. After 1 hour, non-adherent cells were removed, and the medium was refreshed on days 3–4. Adherent cells were used as macrophages for assays on days 7-8, by which time 1-3% of PBMCs had differentiated. These macrophages (5 × 10□ cells/well) were then seeded in 96-well plates and co-cultured with GFP-expressing HLA-A2-positive NALM6 target cells (1 × 10□ cells/well) at an effector-to-target (E:T) ratio of 1:2. The co-cultures were incubated with either isotype control antibody, cAN7, or mAN7. After incubation, phagocytic activity was quantified by flow cytometry, measuring the percentage of macrophages that had internalized GFP-positive target cells.

### LILRB1 binding assay

To evaluate AN7’s capacity to inhibit LILRB1-mediated interactions with HLA class I molecules, a competitive binding assay was performed using recombinant LILRB1-Fc (bio-techne, #2017-T2) and HLA-A2:01-expressed magnetic beads. HLA-A2:01 beads were incubated with 2 µg/mL LILRB1-Fc for 1 hour at room temperature in the presence or absence of 2 µg AN7 monoclonal antibody. For the experimental condition, AN7 was pre-incubated with HLA-A2:01 beads for 30 minutes at 37°C before LILRB1-Fc exposure. Bound LILRB1-Fc was detected using a phycoerythrin-conjugated anti-human Fc secondary antibody and quantified via flow cytometry.

### GVHD mouse models

To assess the therapeutic efficacy of AN7, two distinct preclinical models of graft-versus-host disease (GVHD) were employed.

#### Xenogeneic GVHD Model

The xenogeneic GVHD model was adapted from Nakauchi et al. NSG mice received an intravenous injection of 10 × 10□ HLA-A2□human peripheral blood mononuclear cells (PBMCs) on day 0. This approach induces rapid expansion of human T cells and xenogeneic tissue targeting, resulting in lethal GVHD within 2–3 weeks. On day 2 post-transplant, mice were administered a single intravenous dose of AN7 (100 μg) or an IgG isotype control.

#### Allogeneic GVHD Model

To overcome the limitation of absent hematopoietic stem cell (HSC) engraftment in the xenogeneic system, an immunocompetent allogeneic GVHD model was established. BALB/c mice were lethally irradiated (9.5 Gy) and transplanted with 5 × 10□bone marrow cells from HLA-A2 transgenic C57BL/6 donors. This fully MHC-mismatched combination induces rapidly fatal acute GVHD while permitting donor HSC engraftment. AN7 was administered intravenously at doses ranging from 0.1 ng to 60 μg on day 7 post-transplant. Survival was monitored for 60 days. This model allowed for precise evaluation of allele-specific interventions while preserving critical host-donor immune interactions.

### Mixed Lymphocyte Reaction (MLR) assay

Spleen cells were isolated from HLA-A2 transgenic (Tg) mice (responder cells, designated _RCs_) and BALB/c mice (stimulator cells, designated _SCs_). Bone marrow (BM) cells were thawed from frozen stocks. SCs were pretreated with 25 µg/mL mitomycin C (MMC) for 30 minutes at 37°C. RCs and SCs were co-cultured in triplicate wells of a 96-well plate at a 1:1 ratio (5 × 10□ cells/well each) in RPMI-1640 medium supplemented with 10% fetal bovine serum (FBS) and 1% penicillin/streptomycin. Six antibody conditions were tested: three concentrations (0.001, 0.1, and 10.0 ng) of the experimental antibody (AN7) and matched concentrations of an isotype control (whole mouse IgG). Following a 72-hour incubation at 37°C with 5% CO□, cells were stained with fluorochrome-conjugated antibodies: APC-Cy7 anti-mouse CD3 (clone 17A2), BUV805 anti-mouse CD4 (clone HIB19), BUV661 anti-mouse CD8a (clone 5H10-1), and PE anti-HLA-ABC (clone W6/32). According to the manufacturer’s protocol, absolute cell counts were determined using Count Bright Beads (Invitrogen, #C36950).

### Statics

For all *in vitro* and *in vivo* experiments, Prism9 (Graph-Pad Software) was used to perform statistical analysis. A paired/unpaired t test was used to define statistical significance (*, P < 0.05;**, P < 0.01; ***, P < 0.001; ****, P < 0.0001). One-way ANOVA tests were performed for experiments with more than two conditions. Unless otherwise noted, experiments were performed with at least three biological replicates, with technical duplicates or triplicates per biological sample.

## Acknowledgments

We are deeply grateful to Amy Fan and Asiri Ediriwickrema for their insightful critiques and invaluable guidance, which significantly advanced the quality and direction of this work. We also wish to thank Feifei Zhao for her exceptional dedication and technical expertise in the laboratory, without which this study would not have been possible. This work was mentored and financially supported by Stanford’s SPARK Translational Research Program. We would like to thank all the mentors at SPARK, especially Dirk Mendel, PhD, Kevin Grimes, MD, and Daria Mochly-Rosen, PhD, who believed in and supported our work from start to finish. This work was supported by the Stanford Clinical and Translational Science Award (CTSA) to Spectrum (UL1TR003142). The CTSA program is led by the National Center for Advancing Translational Sciences (NCATS) at the National Institutes of Health (NIH). The content is solely the responsibility of the authors and does not necessarily represent the official views of the NIH. We gratefully acknowledge the financial support for this research provided by SPARK at Stanford. SPARK is supported by Bio-X, Stanford’s interdisciplinary biosciences institute, the NIH Clinical & Translational Science Award (CTSA) through the Stanford Center for Clinical and Translational Research and Education (Spectrum), and the Weston Havens Foundation. We acknowledge Alyssa Chang’s support from a California Institute for Regenerative Medicine (CIRM) Research Training Fellowship.

## Author contributions and disclosures

KN and YN conceived the project, designed and performed the experiments, supervised the experiments, analyzed the data, prepared the figures and tables, and wrote the manuscript. KN, AHC, RM, HN, and YN contributed to the writing of the article. MM, GB, JSK, EM, AB, and SB provided critical clinical opinions. All authors reviewed the final version of the manuscript.

## Refference

1. Smallbone P, Mehta RS, Alousi A. Steroid Refractory Acute GVHD: The Hope for a Better Tomorrow. Am J Hematol. May 2025;100 Suppl :14–29. doi:10.1002/ajh.27592

2. Meyers G, Hamadani M, Martens M, et al. Lessons learned from early closure of a clinical trial for steroid-refractory acute GVHD. Bone Marrow Transplant. Feb 2022;57(2):302–303. doi:10.1038/s41409-021-01529-x

3. Jamy O, Zeiser R, Chen YB. Novel developments in the prophylaxis and treatment of acute GVHD. Blood. Sep 21 2023;142(12):1037–1046. doi:10.1182/blood.2023020073

4. Malard F, Holler E, Sandmaier BM, Huang H, Mohty M. Acute graft-versus-host disease. Nat Rev Dis Primers. Jun 08 2023;9(1):27. doi:10.1038/s41572-023-00438-1

5. Kneifel F, Vogel T, Bormann E, et al. Graft-versus-host disease following liver transplantation: A systematic review of literature. Hepatol Commun. Oct 01 2023;7(10) doi:10.1097/HC9.0000000000000260

6. Cooper JP, Abkowitz JL. How I diagnose and treat acute graft-versus-host disease after solid organ transplantation. Blood. Mar 09 2023;141(10):1136–1146. doi:10.1182/blood.2022015954

7. Nakauchi Y, Yamazaki S, Napier SC, et al. Effective treatment against severe graft-versus-host disease with allele-specific anti-HLA monoclonal antibody in a humanized mouse model. Exp Hematol. Feb 2015;43(2):79–88.e1-4. doi:10.1016/j.exphem.2014.10.008

8. Zuber J, Boyer O, Neven B, et al. Donor-targeted serotherapy as a rescue therapy for steroid-resistant acute GVHD after HLA-mismatched kidney transplantation. Am J Transplant. Aug 2020;20(8):2243–2253. doi:10.1111/ajt.15827

9. Yamazaki S, Suzuki N, Saito T, et al. A rapid and efficient strategy to generate allele-specific anti-HLA monoclonal antibodies. J Immunol Methods. Mar 31 2009;343(1):56–60. doi:10.1016/j.jim.2009.01.007

10. Niu L, Cheng H, Zhang S, et al. Structural basis for the differential classification of HLA-A*6802 and HLA-A*6801 into the A2 and A3 supertypes. Mol Immunol. Oct 2013;55(3-4):381–92. doi:10.1016/j.molimm.2013.03.015

11. Barkal AA, Weiskopf K, Kao KS, et al. Engagement of MHC class I by the inhibitory receptor LILRB1 suppresses macrophages and is a target of cancer immunotherapy. Nat Immunol. Jan 2018;19(1):76–84. doi:10.1038/s41590-017-0004-z

12. Ali N, Flutter B, Sanchez Rodriguez R, et al. Xenogeneic graft-versus-host-disease in NOD-scid IL-2R_γ_null mice display a T-effector memory phenotype. PLoS One. 2012;7(8):e44219. doi:10.1371/journal.pone.0044219

13. McDaniel Mims B, Furr KL, Enriquez J, Grisham MB. Improving bench-to-bedside translation for acute graft-versus-host disease models. Dis Model Mech. Feb 01 2025;18(2) doi:10.1242/dmm.052084

14. Le AX, Bernhard EJ, Holterman MJ, et al. Cytotoxic T cell responses in HLA-A2.1 transgenic mice. Recognition of HLA alloantigens and utilization of HLA-A2.1 as a restriction element. J Immunol. Feb 15 1989;142(4):1366–71.

15. Essers MA, Offner S, Blanco-Bose WE, et al. IFNalpha activates dormant haematopoietic stem cells in vivo. Nature. Apr 16 2009;458(7240):904–8. doi:10.1038/nature07815

16. Tasian SK, Bornhäuser M, Rutella S. Targeting Leukemia Stem Cells in the Bone Marrow Niche. Biomedicines. Feb 21 2018;6(1) doi:10.3390/biomedicines6010022

17. Cao K, Hollenbach J, Shi X, Shi W, Chopek M, Fernández-Viña MA. Analysis of the frequencies of HLA-A, B, and C alleles and haplotypes in the five major ethnic groups of the United States reveals high levels of diversity in these loci and contrasting distribution patterns in these populations. Hum Immunol. Sep 2001;62(9):1009–30. doi:10.1016/s0198-8859(01)00298-1

18. Gragert L, Madbouly A, Freeman J, Maiers M. Six-locus high resolution HLA haplotype frequencies derived from mixed-resolution DNA typing for the entire US donor registry. Hum Immunol. Oct 2013;74(10):1313–20. doi:10.1016/j.humimm.2013.06.025

